# Non-linear functional brain co-activations in short-term memory distortion tasks

**DOI:** 10.1101/2021.09.12.459960

**Authors:** Anna Ceglarek, Jeremi K. Ochab, Ignacio Cifre, Magdalena Fąfrowicz, Barbara Sikora-Wachowicz, Koryna Lewandowska, Bartosz Bohaterewicz, Tadeusz Marek, Dante R. Chialvo

## Abstract

Recent works shed light on the neural correlates of true and false recognition and the influence of time of day on cognitive performance. The current study aimed to investigate the modulation of the false memory formation by the time of day using a non-linear correlation analysis originally designed for fMRI resting-state data. Fifty-four young and healthy participants (32 females, mean age: 24.17 y.o., SD: 3.56 y.o.) performed in MR scanner the modified Deese-Roediger-McDermott paradigm in short-term memory during one session in the morning and another in the evening. Subjects’ responses were modeled with a general linear model, which includes as a predictor the non-linear correlations of regional BOLD activity with the stimuli, separately for encoding and retrieval phases. The results show the dependence of the non-linear correlations measures with the time of day and the type of the probe. In addition, the results indicate differences in the correlations measures with hippocampal regions between positive and lure probes. Besides confirming previous results on the influence of time-of-day on cognitive performance, the study demonstrates the effectiveness of the non-linear correlation analysis method for the characterization of fMRI task paradigms.

## Introduction

The process of cognitive control supports adaptive responses and inhibits automatic ones. It is linked to the working memory not only by function but also by location in the brain – in the prefrontal cortex (for review, see: Miller, 2000). Cognitive control is also enormously involved in decision-making processes to obtain appropriate responses to changing environmental conditions. A model of simple, two-choice decisions that has gained popularity in recent years is the drift-diffusion model (DDM; Ratcliff, 1978; Ratcliff and McKoon, 2008). It describes the decision-making process as an accumulation of evidence about a stimulus from perceptual organs or memory, leading to a reaction (most often a motor one) when the evidence exceeds a certain threshold. The studies investigating neural underpinnings of decision-making focus mainly on the prefrontal areas but increasingly also on the prefrontal-basal ganglia loop (e.g., Bogacz and Gurney, 2007; Sieveritz et al., 2019). Moreover, the interaction between the basal ganglia and the frontal regions has been proven in working memory access control: the basal ganglia detects the appropriate context for a motor response to a stimulus stored in memory (McNab and Klingberg, 2008; Baier et al., 2010; Guo et al., 2018).

Working memory is currently viewed as a multi-component system consisting of three subsystems and a central executive one (Baddeley, 2003). These subsystems include the visuospatial sketchpad, which enables storage of visual information, the phonological loop involved in auditory and verbal information storage, and the episodic buffer, which integrates information from other components keeping a continuous sequence. For many years, researchers attempted to understand the neural correlates of information remembering and retrieving. The hippocampus is a neural structure whose participation in both long-term and working memory has been confirmed in many studies (e.g., Olson et al., 2006; Yonelinas, 2013; Libby et al., 2014). It is also well known that the human memory is prone to errors (Loftus, 1979; Schacter et al., 1998), a fact that motivates the investigation of memory distortion (i.e., false memories) as a byproduct of the memory system, attempting to reveal their nature and mechanism (for meta-analysis on fMRI studies, see: Kurkela and Dennis, 2016). The prevalent paradigm for studying false memories is the DRM (Deese-Roediger-McDermott) paradigm (Deese, 1959; Roediger and McDermott, 1995). Investigating the neural mechanism of false recognition of verbal or nonverbal stimuli is relevant for both encoding (Kim and Cabeza, 2006) and retrieval phases (Schacter et al., 1998). Regarding the false alarms (i.e., when participants incorrectly claim that a new, similar stimulus has appeared previously), most fMRI studies have been using verbal material (e.g. Kim and Cabeza, 2006; Atkins and Reuter-Lorenz, 2011), while the studies using visual objects are less frequent (Lewandowska et al., 2019; Sikora-Wachowicz et al., 2019, 2021). Neuroimaging studies with pictorial material revealed increased activation in the anterior cingulate cortex (Sikora-Wachowicz et al., 2021) and frontal, parietal, and visual cortices (Slotnick and Schacter, 2004; Garoff-Eaton et al., 2005; Gutchess and Schacter, 2012) related to false recognitions.

According to Borbély’s two-process model, circadian processes, the endogenous oscillatory pacemaker, and the homeostatic sleep pressure, which increases with time spent awake, regulate overall human performance and behavior during the 24h cycle (Borbély, 1982; Daan et al., 1984; Borbély et al., 2016). Indeed a large number of studies consistently revealed differences in the cognitive domain according to the time of day (for review, see: Schmidt et al., 2007). Alertness, attention, executive functions, among many others, can be affected by the circadian clock. Also, both short- and long-term memory might be modulated by the time of day (Fabbri et al., 2013; Schmidt et al., 2015).

The current study aims to find neural correlates of encoding and retrieval and the diurnal activity of those correlates in the modified DRM paradigm with abstract, visual objects using a new analysis method – non-linear correlation implemented to the task environment. A typical approach to establish a functional proxy for brain connectivity is to calculate the Pearson linear correlation between the brain’s blood oxygenation level dependent (BOLD) time series and a given stimulus of interest. The alternative used here is motivated by the fact that BOLD fluctuations of a relatively large amplitude capture most of the information (Tagliazucchi et al., 2011, 2012; Liu and Duyn, 2013; Petridou et al., 2013). In consequence, bursts of correlated activity across the brain may be efficiently described by a point process consisting of few discrete events (Cifre et al., 2020). The result was subsequently observed with related methods (Liu et al. 2013; Allan et al., 2015; Karahanoglu and Van De Ville, 2015), and co-activation patterns driven by the point process were studied also in the task paradigm (Jiang et al. 2014, Chen et al. 2015) and in the clinical context (Li et al. 2014; Amico et al. 2014). Following that work, our correlation analysis focuses on the brief instances of large-amplitude signals (so-called events), a technique that may increase the signal-to-noise ratio significantly. Although this method has been proven effective in analyzing brain resting-state data (Tagliazucchi et al., 2011; Tagliazucchi et al., 2012; Cifre et al., 2021), the present study is the first to implement it for signals extracted during a task.

## Materials and Methods

### Participants

As many as 5354 volunteers applied to the first selection stage through the lab’s website announcements. All of them were asked to complete a sleep-wake online assessment including diurnal preference – Chronotype Questionnaire (Oginska et al., 2017), night sleep quality – Pittsburgh Sleep Quality Index (PSQI) (Buysse et al., 1989), and daytime sleepiness – Epworth Sleepiness Scale (ESS) (Johns, 1991). Based on the Chronotype Questionnaire, the individuals were divided into morning and evening chronotypes. Then, 451 participants were qualified for the next selection step included genetic testing for the polymorphism of clock gene PER3, which has been established as a hallmark of extreme diurnal preferences (Archer et al., 2003). After selection, fifty-four volunteers participated in the analysis (32 females, mean age: 24.17 ± 3.56 y.o.) divided into 26 morning types (mean age: 24.31 ± 3.74 y.o.) and 28 evening types (mean age: 24.04 ± 3.24 y.o.). Exclusion criteria were: age below 19 and above 35, left-handedness (assessed by the Edinburgh Handedness Inventory), psychiatric or neurological disorder, drug, alcohol, or nicotine dependence, shift work or travel involving moving between more than two time zones within the past two months, and sleep problems (a result above 10 points from ESS caused exclusion). Subjects did not have any contraindications for magnetic resonance imaging studies. The volunteers were remunerated for participation in the experiment. Prior to the completion of study procedures, they were asked to sign a consent form. The study was conducted under the Declaration of Helsinki and approved by the Research Ethics Committee at the Institute of Applied Psychology at the Jagiellonian University, Krakow, Poland.

### Task

The tasks procedure was based on the DRM paradigm established to investigate the false memories in long-term memory. Given differences between the two types (long- and short-term), the modified version to study short-term memory was developed (Atkins and Reuter-Lorenz, 2011). Two tasks using non-verbal material (abstract, visual objects) requiring global and local information processing were analyzed. The participants had to memorize the set of two stimuli followed by a mask. Subsequently, the stimulus (probe) was displayed, for which a reaction was required, whether the stimulus presented on the screen was present in the preceding set (“yes” with the right hand, “no” with the left). There were three conditions: positive probe (in which the stimulus had been presented in the preceding set), negative probe (the probe had not been presented earlier), and lure (in which the stimulus was very perceptually similar to these in the preceding set but it had not been presented). The third condition seems to produce false memories. Lure probes differed from stimuli in the preceding set in a holistic way (in the ‘global’ task) or individual details (in the ‘local’ task).

There were 60 memory sets presented for 1800 ms followed by 25 positive probes, 25 lures, and 10 negative probes presented for 2000 ms. The memory set and mask were separated by 1000 ms, whereas the mask and probe by 2000-16000 ms (avg. 6097 ms). The two versions of the tasks were created (for morning and evening sessions); each had six versions of the procedure and was counterbalanced within subjects. The dark gray (RGB 72, 72, 72) stimuli were presented on a light-gray background (RGB 176, 176, 176). The abstract objects (5° wide and 4° high) in memory sets were displayed 3° from the screen center to the left and right, while masks and the objects in memory probes in the center of the screen. The task was prepared using E-Prime 2.0 (Psychology Software Tools) and performed during fMRI sessions. The previous study (Ceglarek et al., 2021) describes the task in detail. The example task procedure and analysis flow is depicted in Figure 1.

**Figure 1.**
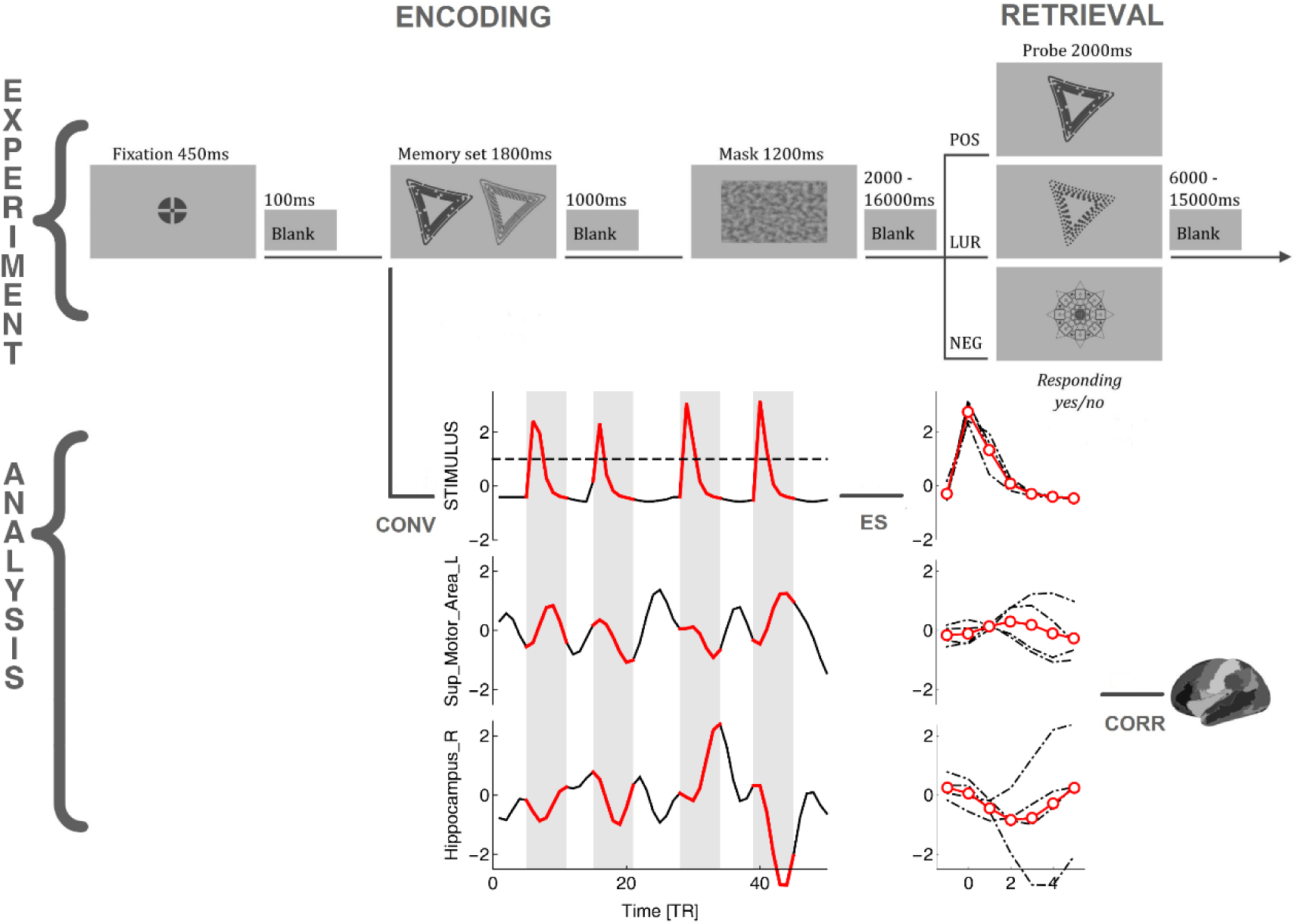
The flowchart of the task requiring global information processing and analysis procedure. POS/LUR/NEG - positive, lure and negative probes, CONV - convolution of the stimuli with a model hemodynamic response function, ES - identification, extraction and averaging of BOLD events, CORR - correlating BOLD events with stimuli, Fisher transforming and performing GLM analysis.

### Procedure

The participants were asked to sleep well (at least 8 hours) the week before and during the entire experimental period. The duration and quality of sleep in the preceding week were controlled using the MotionWatch8 actigraphs (CamNtech, Cambridge, UK). The MR acquisition was conducted twice: in the morning and evening session (one and 10 hours after waking up, respectively; cf. Schmidt et al., 2012). The order of sessions was counterbalanced within subjects. Half of the participants started the procedure with the morning session and half with the evening session. The participants spent the night (before or between sessions) in the Malopolska Centre of Biotechnology in Krakow, Poland, in the same building as the fMRI laboratory. Before the proper experiment in the scanner, computer training was conducted to familiarize them with the MR environment and the task. Morning-type participants performed the task between 09:25 AM and 09:55 AM (SD: 1 hour 12 minutes) in the morning and between 06:30 PM and 07:02 PM (SD: 1 hour 26 minutes) in the evening. Evening-type participants performed the task between 11:00 AM and 11:30 AM (SD: 1 hour 17 minutes) in the morning and between 08:40 PM and 09:10 PM (SD: 1 hour 07 minutes) in the evening. High variability in the task execution time resulted from four experimental tasks being performed during each session, presented in a semi-random way (for more information about other tasks, see: Lewandowska et al., 2018). Individuals abstained from alcohol (48 h) and caffeine (24 h) before the first session and were banned from caffeine and alcohol intake during the experimental days.

### Behavioral data analysis

Statistical analyses were performed using Statistica v13.3 (StatSoft, Inc., 2012) software. To observe differences in accuracy (proportion of correct responses) and reaction times (RTs) the generalized linear model (GLM) with accuracy and RTs as dependent variables with sex and chronotype as between-subjects factors, and with time-of-day, response types (correct and false responses to positive, lure and negative probe) and task (global, local) as within-subjects factors was performed. Due to the small number of errors for a negative probe, only the correct recognitions were left for further analysis. The significance level was set at p<0.05, multiple-comparison corrected.

### fMRI data acquisition

Structural and functional data were collected on a 3T scanner Skyra (Siemens Magnetom, Erlangen, Germany) in Malopolska Centre of Biotechnology in Krakow, Poland, with a 64-channel head coil. For task, 709 functional image volumes with 34 contiguous interleaved axial slices were collected with a T2*-weighted echo-planar sequence (TR = 1800 ms, TE = 27 ms, flip angle = 75, FOV = 256 mm, bandwidth 1816 Hz/Px, voxel size: 4×4×4 mm). Structural data were acquired for each participant using a T1-weighted MPRAGE sequence for a detailed reconstruction of anatomy with isotropic voxels (1×1×1.1 mm) in a 256 mm field of view (256 × 256 matrix, 192 slices, TR = 2300 ms, TE = 2.98 ms). Stimuli were projected on a screen positioned behind a subject’s head; participants viewed the screen in a 45° mirror fixated on the top of the head coil.

### fMRI preprocessing

The preprocessing was performed using the Statistical Parametric Mapping software package (SPM12, Welcome Department of Imaging Neuroscience, UCL, London, UK; www.fil.ion.ucl.ac.uk/spm/) and DPABI (V4.2; Yan et al., 2016) implemented in MATLAB (Mathworks, Inc., MA, USA). Functional images were slice-time corrected, realigned using rigid body transformation, co-registered, and normalized to the EPI template in Montreal Neurological Institute (MNI) stereotactic space with a voxel resolution 3×3×3 mm. The data were spatially smoothed using a Gaussian kernel of FWHM 4 mm, detrended and the covariates like motion parameters, mean signal, white matter, and CSF were regressed. The signal was then filtered with a 0.01-0.1 Hz filter, and the time series from 90 regions of interest (ROI) of the AAL atlas were extracted (Tzourio-Mazoyer et al., 2002).

### Non-linear directed functional co-activations

As commented in the introduction, we used a method originally designed to study the correlation between brain regions during the brain resting state (i.e., no task). In the classical approach, one estimates the resting state correlation by calculating some kind of sliding-window linear Pearson correlation between pairs of BOLD time series. In contrast, the method introduced by Tagliazucchi et al. (2011, 2012) and subsequent authors (Liu and Duyn, 2013; Petridou et al., 2013; Allan et al., 2015; Karahanoglu and Van De Ville, 2015; Cifre et al., 2020, 2021) relies on detecting for a given source BOLD time series the relatively high amplitude activity (“events”) and correlating only these epochs with the other target time series, see Figure 1. The approach is naturally connected to the hemodynamic response function (HRF; Wu et al. 2013) and has allowed to demonstrate the correspondence between rest and task BOLD activations (Petridou et al., 2013). Additionally, such a procedure provides the correlations with a straightforward directionality and time-stamps (Cifre et al., 2021). Thanks to these features, we were able to apply the method to the task setting with the series of task stimuli serving as a synthetic source time series, as described below.

### Definition of source and target events

First, as depicted in Figure 1, the times of stimuli appearance were determined for the Memory sets (encoding phase) and Probes (retrieval phase). Next, to generate a time series encoding the stimuli appearance, we created a binary variable with the sampling rate of 1 kHz (higher than the accuracy of stimulus timing measurements from E-Prime 2.0). Then this time series was convolved with the model HRF obtained from SPM12. After that, it was resampled to 1/TR to align it with the actual BOLD signal, and the resulting time series (termed here stimuli time series) was normalized by its standard deviation (i.e., z-scored). In the present approach, the correlation is computed between the “source” and the “target” events. The “source” events were extracted from the stimuli time series, as segments of 5 TRs after the signal crossed the threshold of 1, including the crossing itself, which is enough to represent the HRF’s whole positive peak. Finally, the “target” events were the segments extracted from the BOLD time series of each of the 90 AAL ROIs at precisely the same times as the “source” events.

### Correlations

The linear Pearson correlation between source and target events was computed and averaged for each experimental condition (subject, session time, phase, probe type, response, and ROI), and the averages were Fisher transformed. The values of these correlations indicate whether a particular ROI systematically co-activated (positive correlations) or deactivated (negative correlations) with a given stimulus type in a given condition.

### fMRI data analysis

Statistical analyses were performed using the R *stats* package (R Core Team, 2021) and the estimated marginal mean package *emmeans* (Lenth, 2021); the data and R scripts are provided in the Supplementary Materials. The general linear model (GLM) assumed *response type* (“yes,” “no”) as the predicted variable, the *phase* (retrieval, encoding), *probe* (positive, lure; the negative probe was not used due to the small number of errors, resulting in no predictive value), *condition* (morning, evening) and *ROI* (90 AAL regions) as the nominal predictors, and the average *correlations* as the numerical predictor. Consequently, we used logistic regression with up to four-way interactions *correlation × ROI ×* (all pairs in the set: *phase, probe, condition*), but *a priori* excluding the terms: *phase, ROI* and their interactions with *probe* and *condition*, since responses are independent of their levels. Such a model was further reduced by a single term deletion based on the Akaike information criterion. The significance level was set at p<0.05, multiple-comparison corrected (Sidak adjustment). The presented results are estimates transformed back from logit to the original variables.

## Results

### Behavioral data

The GLM with accuracy and RTs as dependent variables with sex and chronotype as between-subjects factors, and with time-of-day, response types (correct and false responses to positive, lure and negative probe) and task (global, local) as within-subjects factors revealed significant influence of response type (F(1,8)=439; p=0.001; η^2^_p_=0.64) and interactions: *probe* × *task* (F(1,8)=21,47; p<0.001; η^2^_p_=0.08) and sex × *chronotype* × *probe* (F(1,8)=2.70; p=0.006; η^2^_p_=0.01). The post-hoc HSD Tukey test for accuracy indicated the differences between all response types; for RTs – also between all response types except correct recognition of positive probe vs. correct rejection of lure probe and false responses for positive and lure probes. The descriptive statistics on the proportion of responses and reaction times are presented in Table 1.

**Table 1.** The proportion of responses and reaction times for response types in both tasks.

| probe type | response type | proportion of responses | reaction times (ms) |
| --- | --- | --- | --- |
| | | mean ± SD | mean ± SD |
| positive | hits | 0.82 ± 0.13 | 1277 ± 226 |
| | misses | 0.18 ± 0.12 | 1452 ± 380 |
| lure | correct rejections | 0.74 ± 0.19 | 1295 ± 201 |
| | false alarms | 0.25 ± 0.17 | 1392 ± 419 |
| negative | correct rejections | 0.98 ± 0.08 | 992 ± 161 |
| | false alarms | 0.02 ± 0.04 | 1350 ± 264 |

### fMRI data analysis

The GLMs predicting *response type* (“yes,” “no”) based on *condition, phase, probe type, correlation*, and *ROI* were fitted separately for the global and local processing tasks. The complete type III ANOVA tables for each model are in the Supplementary Materials. Below we report in detail only the highest order significant interactions.

### Task requiring global information processing

The model revealed significant interactions: *condition × correlation* (χ^2^ = 5.26, df = 1, p = 0.022) and *correlation × phase × probe × ROI* (χ^2^ = 332.36, df = 89, p < 2.2*×*10^−16^). In the first case, the contrast between trends of *response* as a function of *correlation* in the *evening* and *morning* conditions (averaged over all *phases, probes*, and *ROIs*) was estimated to be 0.014 (p = 0.022), with the small effect of *correlation* increasing the chance of saying “no” in the evening and of saying “yes” in the morning (*morning* 95% CI [-0.014, 0.0032] and *evening* 95% CI [0.00013, 0.017]), see Figure 2.

**Figure 2.**
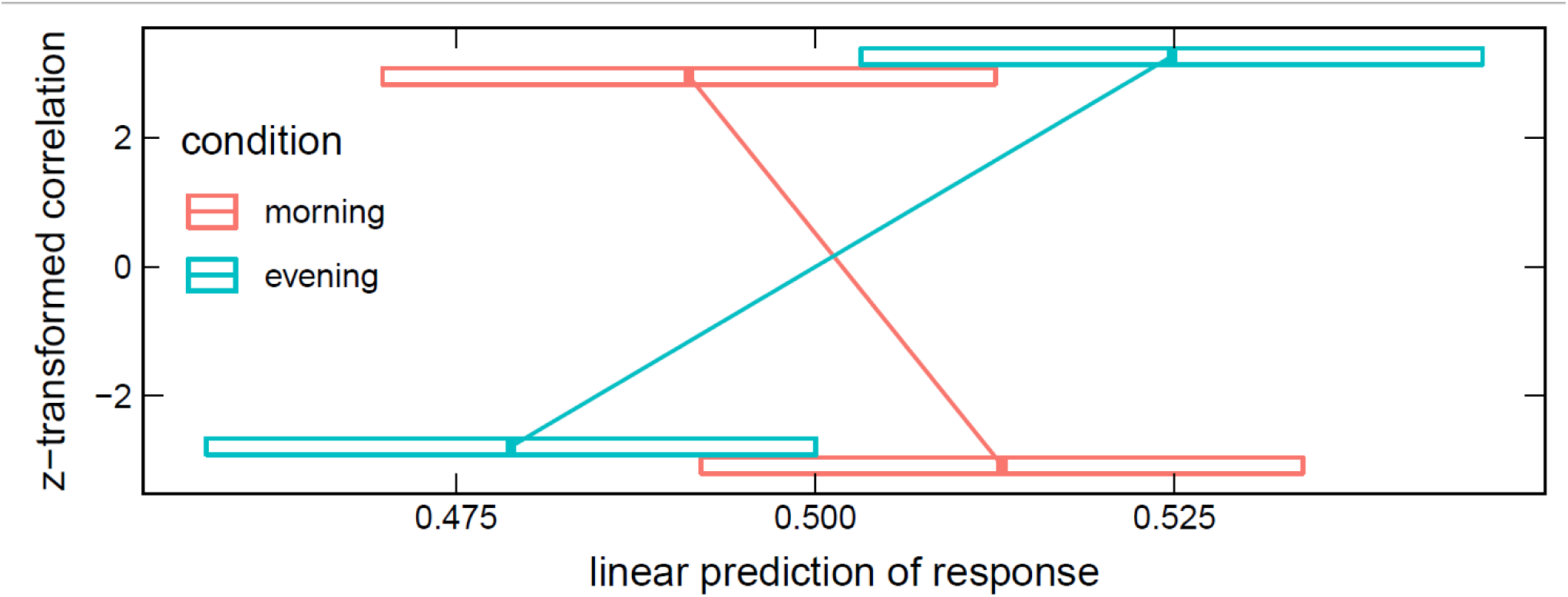
Estimated means’ interaction *condition × correlation* in the global processing task, averaged over *phase, probe* and *ROI* levels. Horizontal bars are 95% CIs.

The interaction *correlation × phase × probe × ROI* disclosed several significant ROIs in encoding and retrieval phases (effects presented in Table 2 and Figure 3). The increasing correlation of hippocampal areas with retrieval stimulus predicted more “no” responses in the *positive* probe and fewer in the *lure*. A reverse effect was observed for the left supplementary motor area (retrieval), olfactory, and medial orbitofrontal cortex (encoding). These brain regions are displayed in Figure 5a.

**Table 2.**
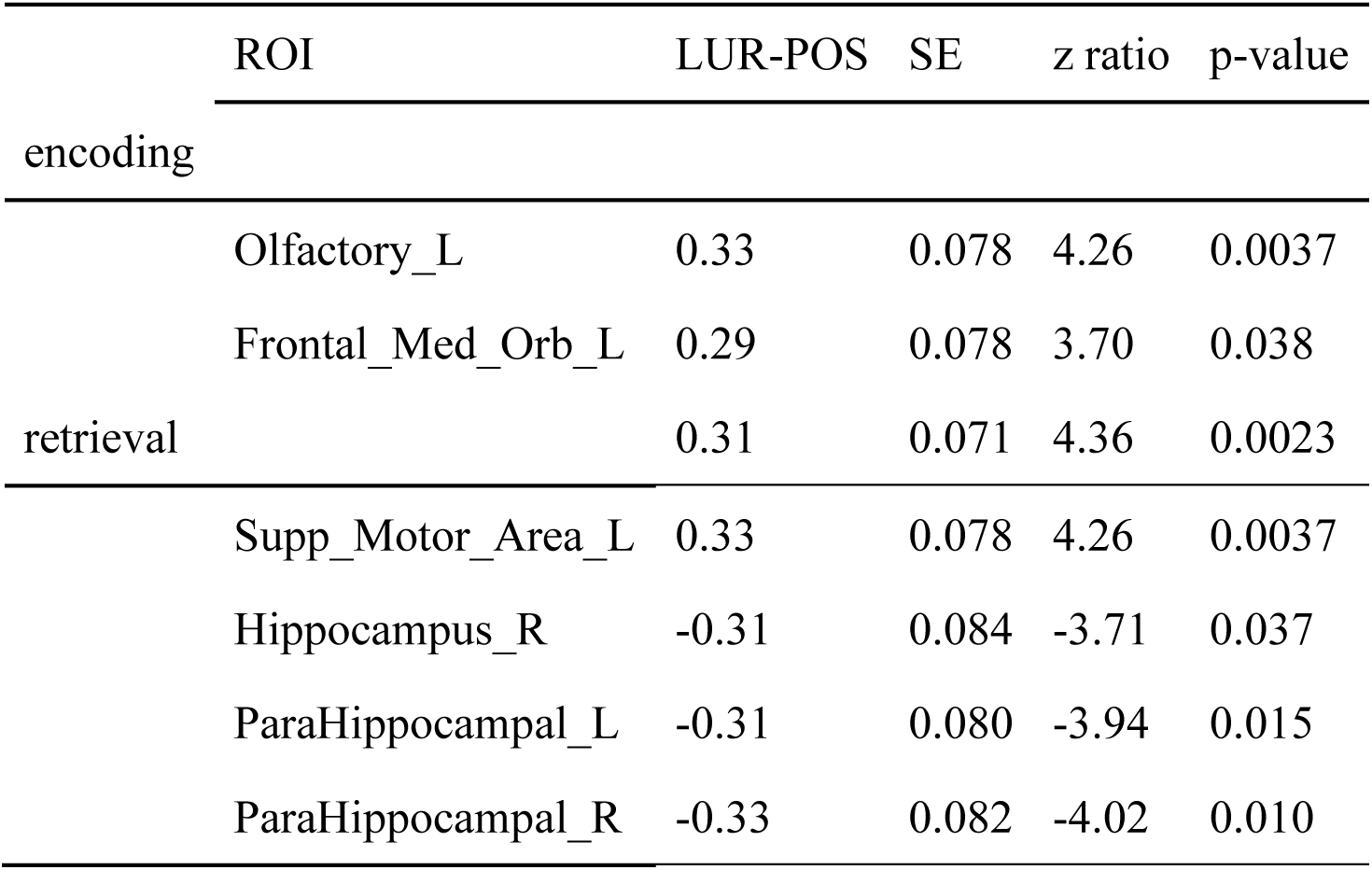
Estimated means’ interaction correlation × phase × probe × ROI in the global processing task, averaged over condition levels. We present the ROIs, where the contrast lure - positive yielded p < 0.05; see Figure 3. Results of all the ROIs are plotted in Figure S1.

**Figure 3.**
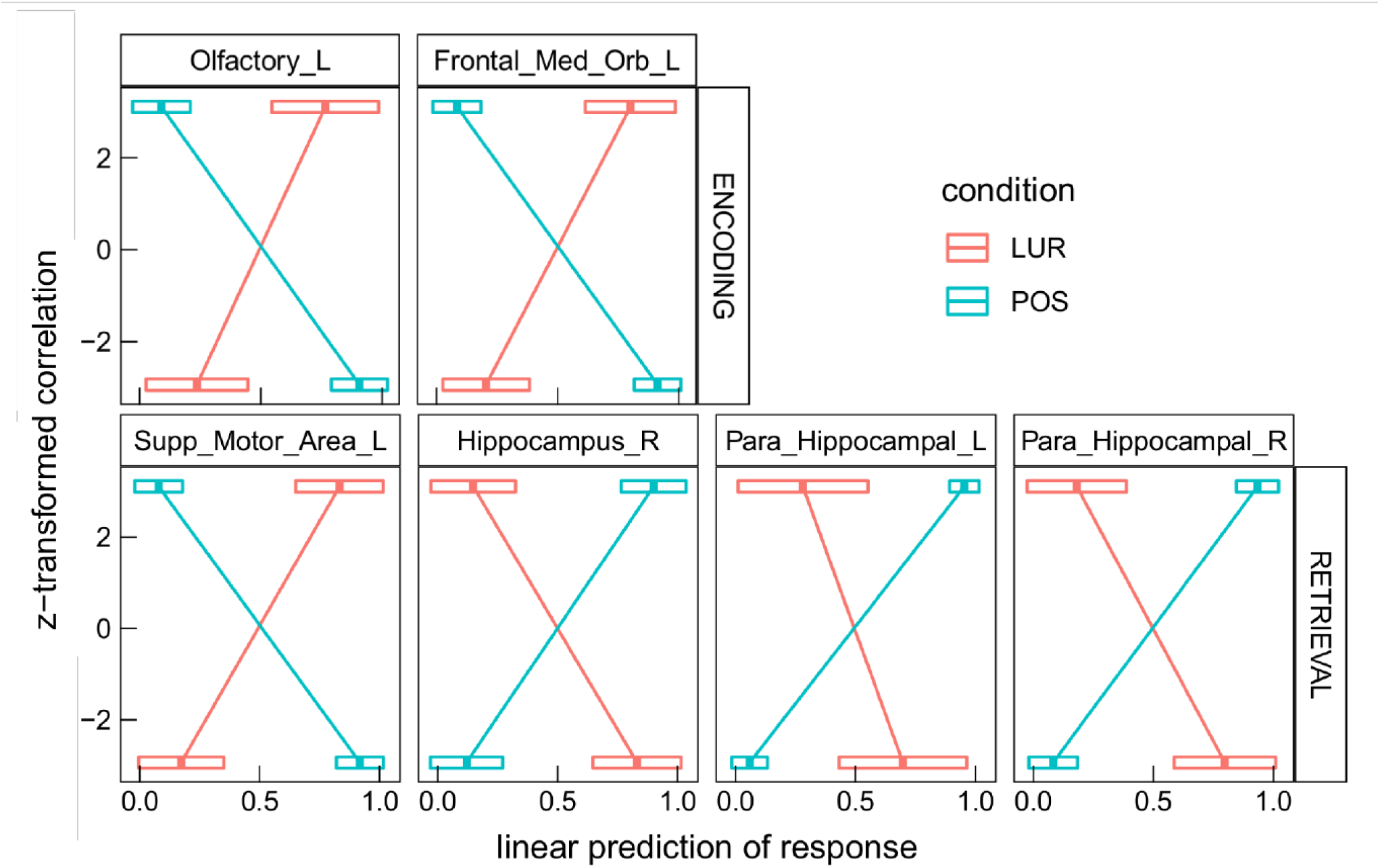
Results of the *correlation × phase × probe × ROI* interaction in the global processing task, averaged over *condition* levels. Horizontal bars are 95% CIs.

### Task requiring local information processing

In the local processing task, the highest order significant interaction containing *condition* was: *condition × probe × correlation × phase* (χ2 = 9.90, df = 1, p = 0.0017). Similarly to the global processing task, the interaction *correlation × phase × probe × ROI* (χ2 = 994.45, df = 89, p < 2.2*×*10^−16^) was also found significant.

In the interaction *condition × probe × correlation × phase* the contrast *morning-evening* was revealed significant for *positive* probe in the encoding phase, 95% CI [0.017, 0.070] (p = 0.0051), and for *lure* probe in the retrieval phase, 95% CI [0.026, 0.078] (p = 0.00043), as well as the contrast *lure*-*positive* probe in the *evening* for both encoding, 95% CI [-0.085, 0.0024] (p = 0.000037), and retrieval, 95% CI [0.0020, 0.087] (p = 0.000066). These results are depicted in Figure 4.

**Figure 4.**
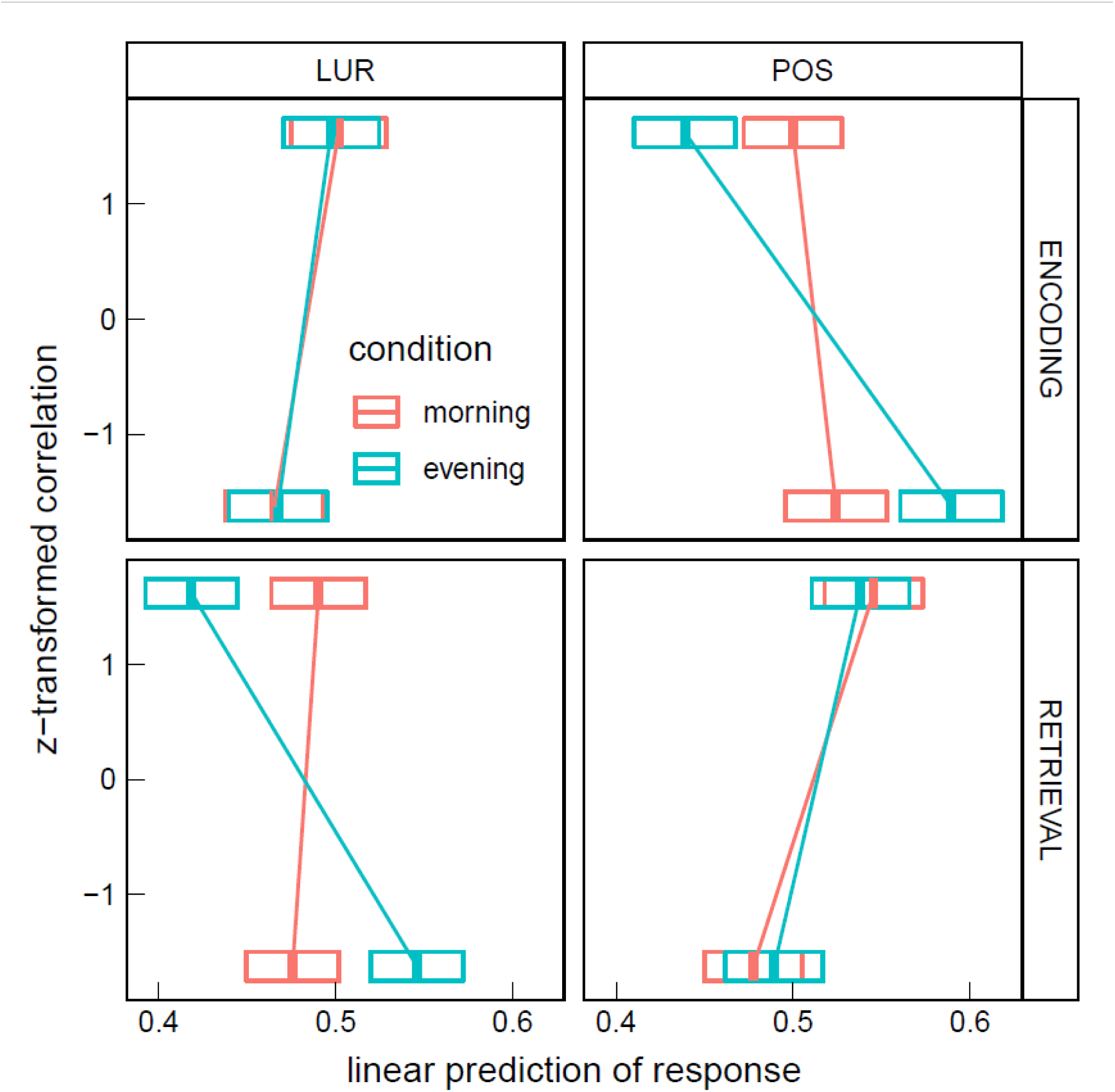
Estimated means’ interaction *condition × probe × correlation × phase* in the local processing task, averaged over *condition* levels. Horizontal bars are 95% CIs.

In the interaction *correlation × phase × probe × ROI*, 18 significant ROIs in encoding and 17 in retrieval phases were disclosed (Table 3). The six most significant brain regions for the encoding (left and right anterior cingulate cortex, left precentral gyrus, left superior frontal gyrus, left superior frontal gyrus medial part, right medial frontal gyrus orbital part) and the retrieval phase (left and right supplementary motor area, right hippocampus, superior temporal pole and left inferior frontal gyrus) are displayed in Figure 5b.

**Table 3.** Results of the correlation × phase × probe × ROI interaction in the local processing task, averaged over condition levels. We present the ROIs, where the contrast lure - positive yielded p<0.05. Six regions having the largest effect correlated with encoding and retrieval stimulus are in bold, and are presented in Figure 5b-c.

| ROI | LUR-POS | SE | z ratio | p-value |
| --- | --- | --- | --- | --- |
| encoding |  |  |  |  |
| **Precentral_L** | -0.45 | 0.095 | -4.7 | 0.00045 |
| Precentral_R | -0.34 | 0.092 | -3.7 | 0.032 |
| **Frontal_Sup_L** | 0.45 | 0.085 | 5.3 | 1.70E-05 |
| Frontal_Sup_R | 0.39 | 0.094 | 4.1 | 0.0072 |
| **Frontal_Sup_Medial_L** | 0.43 | 0.082 | 5.3 | 2.50E-05 |
| Frontal_Sup_Medial_R | 0.37 | 0.087 | 4.3 | 0.0036 |
| Frontal_Med_Orb_L | 0.34 | 0.083 | 4.1 | 0.0077 |
| **Frontal_Med_Orb_R** | 0.39 | 0.085 | 4.6 | 0.00075 |
| **Cingulum_Ant_L** | 0.41 | 0.084 | 4.9 | 0.00014 |
| **Cingulum_Ant_R** | 0.52 | 0.095 | 5.4 | 9.10E-06 |
| Cingulum_Post_L | 0.39 | 0.08 | 4.8 | 0.00026 |
| Cingulum_Post_R | 0.34 | 0.082 | 4.2 | 0.0046 |
| Para_Hippocampal_R | 0.35 | 0.093 | 3.8 | 0.030 |
| Parietal_Inf_L | -0.35 | 0.084 | -4.2 | 0.0058 |
| Parietal_Inf_R | -0.36 | 0.089 | -4.1 | 0.0082 |
| Temporal_Pole_Sup_R | 0.38 | 0.093 | 4.1 | 0.0074 |
| Temporal_Mid_R | 0.34 | 0.091 | 3.7 | 0.041 |
| Temporal_Pole_Mid_R | 0.38 | 0.091 | 4.1 | 0.0067 |
| retrieval |  |  |  |  |
| Precentral_L | 0.42 | 0.087 | 4.8 | 0.00027 |
| Frontal_Mid_L | 0.41 | 0.093 | 4.4 | 0.0017 |
| Frontal_Mid_Orb_L | 0.32 | 0.085 | 3.8 | 0.027 |
| **Frontal_Inf_Oper_L** | 0.5 | 0.097 | 5.2 | 3.90E-05 |
| **Frontal_Inf_Oper_R** | 0.49 | 0.11 | 4.4 | 0.0016 |
| **Supp_Motor_Area_L** | 0.57 | 0.09 | 6.3 | 6.00E-08 |
| **Supp_Motor_Area_R** | 0.54 | 0.1 | 5.3 | 2.20E-05 |
| Hippocampus_L | -0.38 | 0.093 | -4.1 | 0.0065 |
| **Hippocampus_R** | -0.51 | 0.097 | -5.3 | 2.60E-05 |
| Para_Hippocampal_R | -0.44 | 0.087 | -5 | 9.00E-05 |
| Amygdala_R | -0.39 | 0.087 | -4.5 | 0.0013 |
| Cuneus_L | -0.37 | 0.087 | -4.3 | 0.0036 |
| Parietal_Sup_L | 0.42 | 0.095 | 4.4 | 0.0018 |
| Temporal_Sup_L | -0.39 | 0.095 | -4.1 | 0.0089 |
| Temporal_Pole_Sup_L | -0.36 | 0.092 | -4 | 0.012 |
| **Temporal_Pole_Sup_R** | -0.53 | 0.098 | -5.4 | 9.20E-06 |
| Temporal_Pole_Mid_R | -0.34 | 0.09 | -3.7 | 0.032 |

**Figure 5.**
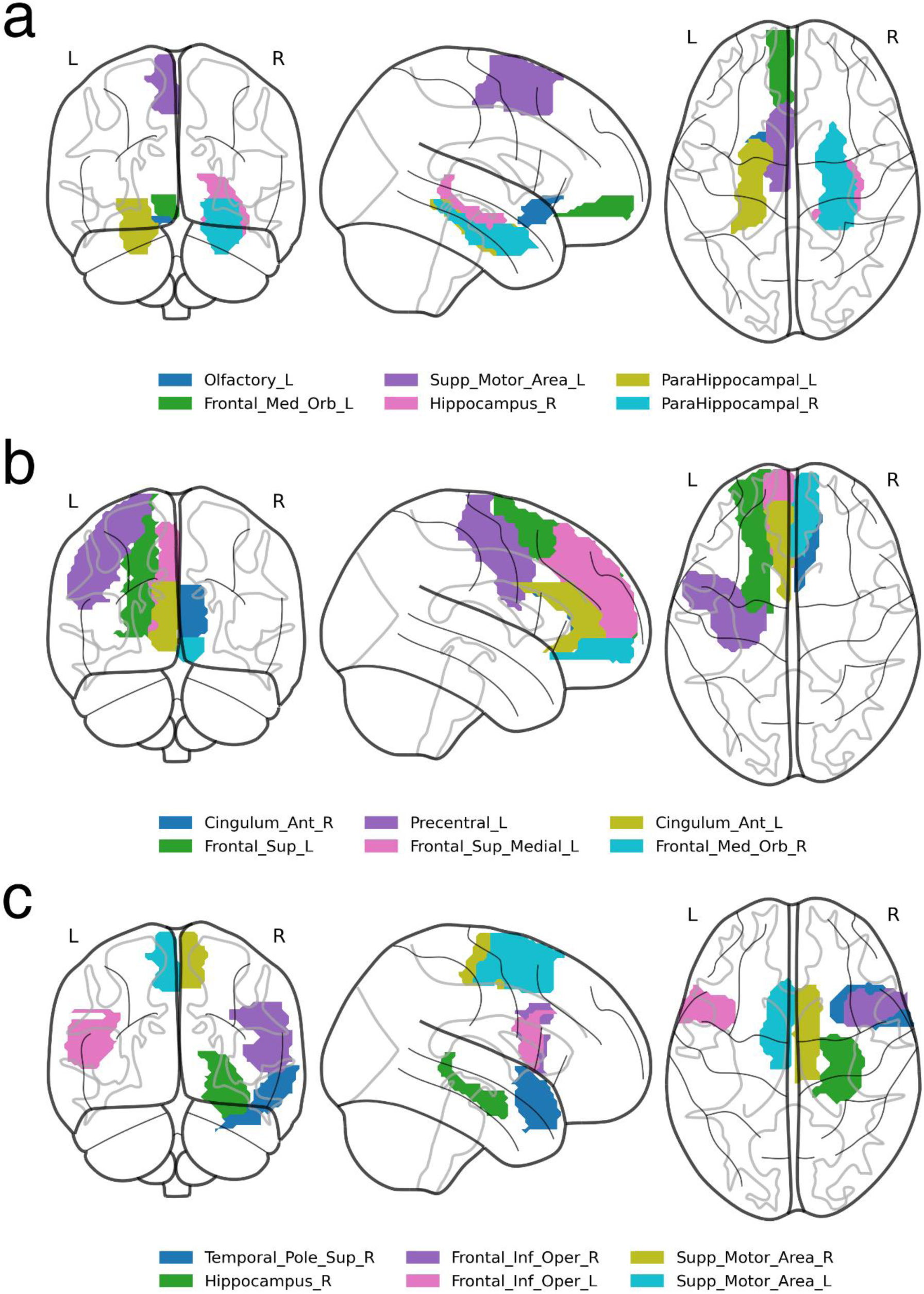
Visualization of brain regions whose correlation with the stimuli predicts difference in response between *lure* and *positive* probes in: a – global information processing task, b – encoding in local information processing task, c – retrieval in local information processing task.

## Discussion

The current study investigated false memory formation in short-term memory. Employing a novel approach, it revealed features of the neural mechanism behind memory distortions. Although the method has been applied before to resting-state data, according to our knowledge, the present results are the first attempt to employ this technique to task fMRI data. We successfully demonstrate the use of the non-linear correlation method on short-term memory tasks requiring global and local visual processing as well as search for diurnal differences in correlations.

These results indicate some time-of-day effects on the neural mechanism of false memories formation. Previous studies revealed time-of-day differences in neural activity during cognitive tasks (for a review, see: Gaggioni et al., 2014). Marek et al. (2010) confirmed diurnal variations in neural activity of orienting the attentional system during a Stroop-like task. Schmidt and colleagues (2015), using the n-back paradigm with different cognitive load, demonstrated decreased activity in the ventrolateral prefrontal cortex and premotor areas from the morning to the evening hours for the higher cognitive load. The interaction of time-of-day with BOLD correlations that we observe are in general weaker than other ones (e.g., *correlation × probe*). In the global processing task, the overall correlations with stimuli predict a higher proportion of “no” responses in the evening and lower in the morning. However, other interactions (with phase and probe), which appeared significant only in the analysis of the local processing task, might confound this effect. Nevertheless, our result with diurnal variation of responses is consistent with the recent work of Tandoc and colleagues (2021), which revealed an increased generalization process that leads to increased false memory formation in the morning – an effect explained by lower inhibition at morning hours. Another study on resting-state data suggested the less efficient brain networks organization in the first hours after waking, which could be an effect of sleep inertia (Farahani et al., 2021). The stronger effect for local processing, visible in Figure 4, reads that in the evening the increased correlation of the whole brain’s activity with encoding stimulus predicts a lower proportion of “no” responses to a positive probe (more correct responses), and similarly, the increased correlation with retrieval stimulus predicts a lower proportion of “no” responses to a lure probe (more incorrect responses).

Regions that showed significant differences in correlations in both tasks overlapped in the retrieval phase (see: Table 2 and 3), which strengthens the result and demonstrates the effectiveness of the new method of analysis. Regarding the task requiring global information processing, we observed differences in correlations in the orbitofrontal region for the memorizing phase of the task and hippocampal and parahippocampal areas for the retrieval phase. The orbitofrontal cortex is involved in the process of decision making (Steiner and Redish, 2012) and encoding the new visual stimuli (Frey and Petrides, 2000; O’Doherty, 2007). The middle temporal cortex, which includes the hippocampus and parahippocampal gyrus, plays a crucial role in remembering and retrieving events, facts, and details. Moreover, using auditory verbal material, Cabeza et al. (2001) suggested that the parahippocampal gyrus can distinguish between false and true recognition. Garoff-Eaton et al. (2006) confirmed that capacity of the right parahippocampal gyrus with a procedure that uses abstract visual stimuli. The current study supports these results, showing that correlations of the right hippocampus and parahippocampal gyri with the positive and lure probe differentially predict subjects’ responses. Higher correlation in all these areas consistently predicted more “no” responses to positive probe and fewer for lure (both increasing the proportion of incorrect responses), see Figure 3.

There are more differences in correlations in the task requiring local (detailed) information processing, especially in frontal, cingulate, and temporal cortices. For the encoding phase, one sees the activations of the anterior, middle, and posterior cingulate cortex, which are part of the limbic system responsible for regulating emotion, learning, and memory (Rolls, 2019). Extensive psychological studies on the functional organization of the brain revealed the hemispheric functional separation in a way that the left hemisphere is engaged in language processing and the right is responsible for visuospatial functions (e.g., Gazzaniga et al. 1965; Milner 1971; Corballis et al. 1999, 2002; Zuanazzi and Cattaneo 2017). Our task employing abstract objects located in space seems to engage the right hemisphere more, which can be seen in differences in correlations in right temporal cortices.

It is worth mentioning that the type of correlation analysis used here is appealing, given the recent results indicating that BOLD infraslow signal fluctuations throughout the brain are coherent with arousal fluctuations (Raut et al., 2021). These results on humans fMRI show that ongoing arousal fluctuations are correlated with global waves of activity slowly propagating in parallel through the neocortex, thalamus, striatum, and cerebellum. We could speculate that the observed differences between time-of-day may be causally related with these high amplitude waves, corresponding with different degrees of arousal and thus of cognitive performance, a point that deserves further consideration.

This study was the first attempt at implementing the new non-linear coactivations method in the task environment. The limitation of the current application of the method is the possible delay of the BOLD response to the stimuli with respect to the timing of the stimuli themselves. If the dominant peak of the response is 1-5 TRs delayed, then one would observe a negative non-linear correlation with an ROI that, in fact, responds positively to a stimulus, unless the anticipatory vasodilation comes into play (Sirotin and Das, 2009) and synchronizes the BOLD activations. The precise interpretation of an ROI’s role following such statements as “*higher correlation* with an ROI predicts fewer correct responses” is thus contingent on that delay. Incorporating delays into the non-linear correlation analysis is actually possible, as shown in Cifre et al. (2021), and is a topic of our further investigation.

In conclusion, the models generally explained less variance in the global visual-feature processing task data than the local one. The dependence of false memories formation on time-of-day was generally present but weaker than other effects, currently not allowing to pinpoint any particular ROI affected by it. However, diurnal variation of responses could be explained by lower cortical inhibition immediately after sleep, according to the synaptic homeostasis hypothesis. On the other hand, there was enough evidence to find significant differences in processing the positive and the misleading stimuli in specific brain areas. Most notably, we found that peaks in the BOLD signal in the supplementary motor areas immediately after presenting stimuli in the retrieval phase consistently predicted correctness of the following response (“no” for lure and “yes” for positive), and the reverse was true for the hippocampal regions (peaking BOLD at the time of retrieval consistently predicted incorrect responses).

The study demonstrates that non-linear fMRI correlations can be applied effectively to the task paradigm. They were found informative as predictors in generalized linear models, where the interaction terms with atlas-based ROIs indicated specific loci associated with producing responses to the tasks. The method allows finding brain areas related to processing the stimuli and opens new possibilities for analyzing other cognitive tasks.

## Author Contributions

MF, BS-W, KL and TM conceptualized the whole project and established the fMRI tasks procedure, MF, BS-W, KL and BB collected the behavioral and imaging data, AC performed the preprocessing steps, AC and JKO analysed the fMRI data and prepared figures. DRC provided feedback on results interpretation, and AC, JKO, IC and DRC wrote the first draft of the manuscript. All authors reviewed the manuscript.

## Acknowledgments

Work conducted under the auspice of the Jagiellonian University-UNSAM Cooperation Agreement. AC, BB and JKO thanks Dante Chialvo for facilitating their visit and the hospitality of Universidad de San Martín, UNSAM, Argentina. We would like to thank Prof. Patricia Reuter-Lorenz for her constructive suggestions during the planning and development of the project and her valuable support. We also thank Anna Beres and Monika Ostrogorska for their assistance in data collection, Piotr Faba for his technical support on the project and help in data acquisition, and Aleksandra Zyrkowska for help in the process of participants selection.

## Data Availability Statement

The data, scripts and extended results for this study (additional figures) can be found in the OSF repository (https://osf.io/r46wf/).

## Supplementary Materials

The MATLAB code accompanying the paper (Cifre et al., 2021) for computing non-linear functional coactivations is available at https://github.com/remolek/NFC. The data, R scripts, fitted models, and results used in statistical analysis in this paper are available at https://osf.io/r46wf, as well as supplementary figures.

